# Cryogenic Electron Tomography by the Numbers: Charting Underexplored Lineages in Structural Cell Biology

**DOI:** 10.1101/2025.07.09.663989

**Authors:** T. Bertie Ansell, Louis Berrios, Kabir Peay, Peter Dahlberg

## Abstract

Imaging of cells and their interactions across the whole biosphere, with molecular-scale resolution, is key for understanding structure-function relations. Cryogenic electron tomography (cryo-ET) is a powerful method for obtaining this critical information. However, cryo-ET studies are challenging and often limited to a small number of cell types per study. Here, we bring together cryo-ET data from hundreds of cells and tissues across the biosphere to (i) identify emerging methodological trends, (ii) reduce imaging time and costs, (iii) quantitively compare methods for cell freezing and sectioning and (iv) census cryo-ET species coverage across all domains of life. Comparison of the fraction of cellular material within lamellae across all domains of life reveals an order of magnitude difference between bacteria (9%) and archaea (15%) compared to eukaryotes (1%). We calculate the fraction of cellular material which can be imaged using distinct sectioning methods on multicellular communities and tissues - identifying serial lift-out as a powerful new approach for obtaining more complete cellular depictions. Finally, we show that the biodiversity of current cryo-ET studies is 2-3 order of magnitude lower than in sequence libraries and 4-5 lower than the total predicted on Earth. Our analyses reveal major evolutionary lineages which remain critically understudied and where future cryo-ET research would be most impactful.

**Significance Statement:** Cryogenic electron tomography (cryo-ET) is a powerful method for imaging inside cells, but it is challenging and typically limited to a few cells per study. We consolidate and quantify data from hundreds of cellular cryo-ET studies across the biosphere. The methodological trends we identify will enhance cryo-ET throughput and heighten biological insight from underexplored lineages of life on Earth.

## Introduction

Cryogenic electron tomography (cryo-ET) is emerging as the standout method for label-free, non-perturbative imaging of biological material with nanoscale resolution^1^. It is currently the only imaging modality, for instance, which routinely achieves resolutions of 2-4 nm while preserving cells in their native, hydrated states^2,3^. This powerful method has resolved key sub-cellular processes in exquisite detail including (i) *in situ* structures of bacterial flagella^4^ and (ii) injection complexes^5^ which could not be easily purified, (iii) visualisation of cytoskeletal arrangements that enable intracellular trafficking processes^6,7^, (iv) viral capsid structures and infection cycles^8^, (v) the native arrangements and heterogeneity of protein complexes such as ribosomes^9^ and (vi) partitioning of phase-separated regions^10^. The resolution provided by cryo-ET therefore has and continues to yield unparalleled knowledge of biological structures and novel insights of how life functions.

Nonetheless, the cryo-ET pipeline is often cumbersome, with typical timelines of ∼1-2 weeks for culturing cells, freezing grids, sectioning samples and obtaining tomograms. Furthermore, each stage requires highly skilled manual intervention to transfer samples between the different instruments and manipulate them *in situ*. For context, we estimate the total cost of cryo-ET to be ∼US$3-5k for a single imaging round. These high user costs reflect the steep price of instrumentation purchase, maintenance and expert staff, leading to facility operational budgets on the order of millions US$ per year^11^. This is 2-4 orders of magnitude greater than cellular light-microscopy^12^, and is a substantial barrier to cryo-ET research. Resultantly, these constraints tend to limit studies to only one^4^ or a small subset of cell types^13^, which restricts biological insight. The field could therefore benefit from efforts that reduce time and costs and enhance cryo-ET throughput.

One potential solution to these problems is to clarify and consolidate reproducible methods needed to scale cryo-ET applications for imaging across our vast biosphere. These analyses would help (1) identify methodological trends to speed up experimental proceedings, (2) reduce time and labour costs, (3) identify potential paradigm-shifting techniques for moving structural analyses towards more complete depictions of whole cells/organisms, (4) census species coverage within the literature and (5) identify underrepresented areas of life to target in future cryo-ET studies. While there are several reviews which summarise the advances and limitations of cryo-ET^1,14,15^, we are aware of no reports which statistically evaluate cellular cryo-ET studies and which identify factors shaping methodological choice across all domains of life.

To achieve these aims we first elaborate on technicalities of the cryo-ET pipeline necessary for preparation of cells or tissues. Cells must be cultured and frozen on electron microscopy (EM) grids by one of two processes: (i) rapid plunging into liquid ethane under cryogenic temperatures or (ii) contact with liquid nitrogen at ∼2100 bar within a high-pressure freezer (HPF), which depresses the sample freezing point and slows ice crystal growth^16,17^. This results in a layer of vitrified (amorphously frozen) ice typically 1-10 μm (plunge-freezing) or 10-200 μm thick (for HPF). For imaging with the transmission electron microscope (TEM) samples must be <0.3-0.5 μm thick else inelastic scattering of electrons by the cellular material limits the resolution obtainable^18^. Hence, thicker samples must be sectioned, typically using a focussed ion beam (FIB) of accelerated gallium or plasma ions (xenon, argon, oxygen)^19,20^. The FIB beam ablates cellular material with nanometer-scale precision above and below a thin slab known as a lamella. If the sample is extremely thick micro-manipulation techniques may precede FIB milling of lamella, whereby a larger chunk of material is excised using the FIB and deposited on another grid using a needle. Excision of a single block of material is called cryo-FIB lift-out^21^. In serial lift-out 3-4 μm sub-segments of this block are excised and redeposited for independent sectioning^22,23^. An alternative sectioning method involves using an ultramicrotome to precisely cut cellular material using a diamond knife^24^. Finally, once a ∼0.2 μm lamella has been produced, the grid is transferred to the TEM where it is sequentially tilted and imaged to generate a cryo-ET tomogram. Subsequent computational processing and imaging segmentation steps are performed, however, these are not the focus of this study and are detailed elsewhere^25^. Hence, this overview spotlights the considerable expertise necessary for production of a small number of lamellae, which typically account for a tiny subset of the total cellular material. Thus, efforts to consolidate studies and identify trends are essential.

In this study, we consolidated cryo-ET data from hundreds of cell or tissue types across bacterial, archaeal and eukaryotic domains of life. One key goal of this analysis was to identify cellular traits which shape transitions between methodological approaches. We also assess which methods move the field closer to imaging whole cells/tissues – a vital methodological advancement needed for tracing structural architectures across biochemical and spatial gradients. Additionally, we quantitatively map the fraction of cellular material captured within lamellae for distinct cell types and sectioning methods. This is the first comparative statistical analysis of lamellae fractions across the biosphere, strengthening prior estimates of 0.5-4%^26^ and directing future efforts towards increased cell coverage via emerging methods. Finally, we compared the biodiversity within cryo-ET studies to both the sequence diversity in publicly available databases and the total predicted biodiversity on Earth. Our findings expose major evolutionary lineages that remain poorly studied and ripe for cryo-ET exploration.

## Methods

### Literature Search

A literature search of ‘Cryogenic electron tomography’ was performed using the ‘Web of Science’ database and entries were filtered using the ‘Cell Biology’ category (www.webofscience.com). The literature search was performed in August 2024 and updated through December 2024. A total of 210 entries from 1984 to the end of 2024 were identified and filtered further by manual inspection using the following criteria. Results were refined by excluding studies which (1) used resin embedding for sample preparation, (2) did not use intact cells (e.g. visualised proteins, fibrils, cell walls or virus particles) or (3) did not report methods. In addition, reviews, abstracts and thesis dissertations were excluded, as were studies that only performed cryo-TEM. The remaining 53 studies were primary research articles which performed cryo-ET on cells. We subsequently supplemented entries using The Atlas of Bacterial and Archaeal Cell Structure by manually cataloguing articles associated with individual entries within the Jensen Lab cell atlas^27^. This gave an additional 34 entries, bringing the total number of articles to 87.

For papers that reported cryo-ET on multiple cells/tissues (28 studies), repeat entries were created for each cell/tissue type. Hence, a representative sample of 175 cells or tissues are included within this meta-analysis (Supplementary Table 1). Bacterial and archaeal cells are listed according to genus and eukaryotic cells are listed according to cell line or genus (if applicable). There were 100 studies of bacterial, 12 archaeal and 63 eukaryotic cells each.

Each entry was manually inspected and the methods used for freezing and sectioning of cells/tissues were recorded in Supplementary Table 1, including chemical details of cryoprotectants used in a subset of studies (17 entries). This dataset is not an exhaustive list of all cryo-ET studies performed to date since this is not feasible and manual inspection was essential to extract methodological details. However, it is a representative sample of cryo-ET studies, which intends to accurately capture the taxonomic diversity of cells studied.

### Cell/tissue Size Estimates

Cell/tissue sizes (lengths, widths/thicknesses) were mostly derived from the literature. If a range of lengths/widths was given the mean value was used. For entries taken from The Atlas of Bacterial and Archaeal Cell Structure^27^ cellular dimensions were measured directly from images in the database if the entire cell was visible, else derived from the literature (Supplementary Table 1). Each cell was described as either a capsule, sphere, square, rectangle, cone or rectangular pyramid, and the shape dimensions were recorded. For example, the length and width/thickness were reported for capsule, spherical, square and conical cells. For rectangular and rectangular pyramid shaped cells an additional length dimension was recorded.

### Analysis of Imaged verses Total Cellular Fractions

For each sectioning approach (none, FIB-SEM, cryo-FIB lift-out, serial lift-out or ultramicrotome) we calculated the maximum number of lamellae (*Nl*) that could be produced by following a standard sectioning workflow once for each of four commonly studied cells/species (*Escherichia coli, Chlamydomonas reinhardtii*, HeLa and *Caenorhabditis elegans*). For FIB-SEM and cryo-FIB lift-out a single lamella (*Nl* = 1) is produced for each round of sectioning whereas for serial lift-out and the ultramicrotome the cell length (*L*) was divided by the minimum theoretical separation distance between lamellae, assuming a lamellae thickness (*l*) of 0.18 μm^28^. This separation distance was therefore ∼4 μm for serial lift-out^22,23^ or 0.18 μm (i.e. no separation between lamellae) with the ultramicrotome. The percentage of the cell imaged (*P*) was defined using Equation 1.

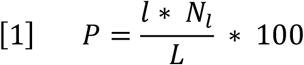

We also tried estimating the fraction of a cell contained within a lamella by comparing lamella volumes to total cellular volumes. However, these estimates were less robust for comparative analyses across domains because lamellae volumes were typically larger than total bacterial and archaeal cell volumes.

### Comparison of Cryo-ET Entries with Global Sequencing Abundance

The number of recorded prokaryotic genera were obtained from the PATRIC database (version 3.49.1, accessed December 2024) based upon the number of entries with rRNA sequencing data (https://www.bv-brc.org/view/Bacteria/2)^29^ and the total predicted genera on Earth are reported in the SILVA database^30,31^ (version 138.2, accessed December 2024). The number of recorded eukaryotic genera were obtained from rRNA entries within EUKARYOME^32^ (version 1.9.4, accessed April 2025) while the total predicted genera were obtained from the Interim Register of Marine and Nonmarine Genera (IRMNG) for extant and accepted eukaryotic entries^33^ (accessed April 2025). The predicted number of eukaryotic species by kingdom were derived from reported taxonomic estimations^34^. The number of human cell lines entries were derived from those listed in the Human Protein Atlas based on RNA sequencing (https://www.proteinatlas.org/humanproteome/cell+line/data#cell_lines)^35,36^. Predicted estimates and database entries across kingdoms are unlikely to be true reflections of species diversity but are necessary approximations for abundances on Earth and useful for comparisons.

### Data Analysis and Summary Reporting

For statistical analyses of the amount of cellular material within lamellae a Kruskal-Wallis test was performed (*p* = 3 × 10^−21^) followed by a pairwise Dunn’s tests between domains (Bact-Euk: *p* = 8 × 10^−19^, Bact.-Arch: *p* = 2 × 10^−1^, Bact-Arch: *p* = 5 × 10^−10^) using the SciPy and Scikit-posthoc python packages^37,38^. The Kruskal-Wallis test was selected because data was not normally distributed and there were different sample sizes across domains. Analysis of data was conducted in python and both the code and associated datasets are deposited on zenodo (DOI: 10.5281/zenodo.15831544). Details of cell studies and publication are provided with the Supplementary Information.

## Results

### The Emerging Cellular Cryo-ET Revolution

To evaluate the current landscape of cellular cryo-ET research, we conducted a meta-analysis of 175 cell or tissue types. Using data pooled from 87 primary research articles released between 2006 and 2024, we consolidated studies of 100 bacterial, 63 eukaryotic and 12 archaeal cells (Supplementary Table 1) to address outstanding information gaps within the literature. Notably to:

- Map methods for freezing and sectioning across a range of cell types.
- Identify factors which shape transition points between methodological approaches.
- Quantitatively assess the fraction of cellular material captured within lamellae.
- Highlight powerful but underutilised methodological approaches.
- Identify under-explored kingdoms of life to expedite future cryo-ET research.

The cryo-EM ‘resolution revolution’ can be attributed to the years starting ∼2012 due to the innovation and commercialisation of direct electron detectors (DEDs) (2004, 2008)^39^, Volta phase plates (2014)^40^, 300 KeV electron microscopes and advances in computational image processing^1,41–43^. Compared to the so-called cryo-EM ‘resolution revolution’, the increase in cellular cryo-ET studies was shifted by approximately 6 years, likely due to the added complexity of *in situ* imaging (Fig. 1). Hence, this systematic analysis of cellular cryo-ET studies is aptly timed to assess the current state of the field and where future efforts would be most impactful.

**Figure 1:**
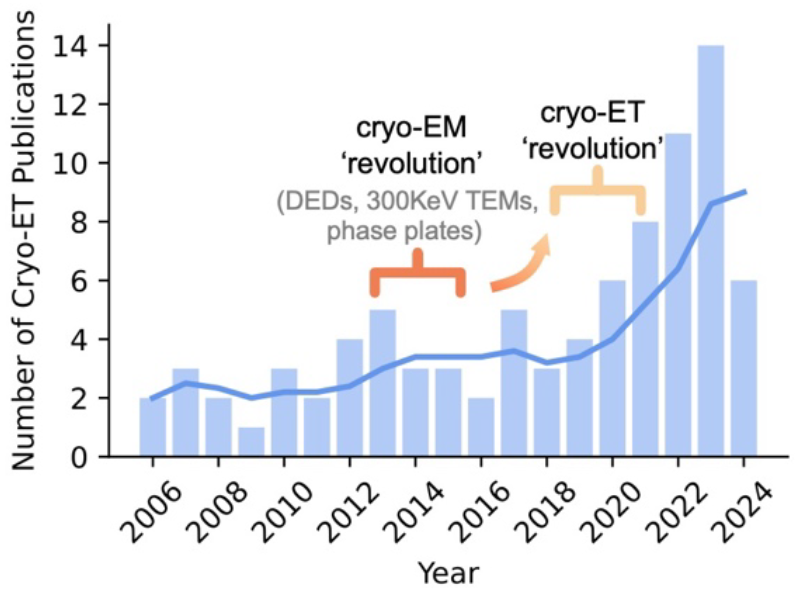
The emergence of a cellular cryo-ET revolution. Summary of the number of filtered publications by year included within this analysis until the end of 2024.

### Cell Size Dictates Choice of Freezing and Sectioning Approach

First, we sought to quantify the extent to which specific cellular traits, such as *size* or cellular *composition*, drive switching between cryo-ET freezing methods^44^. We plotted the length versus width or thickness of cells/tissues and coloured these based on whether plunge freezing or (Fig. 2A, blue) or high-pressure freezing (HPF) (Fig. 2A, red) was used for vitrification. Entries were linearly distributed along cellular dimensional scales (R^2^ = 0.73, *p* = 2.8 × 10^−30^) and ranged from 0.1-0.2 μm at the smallest scales to 100-2000 μm for multicellular material. We found that a transition between plunge-freezing and HPF typically occurred when one cellular dimension surpassed 100 μm, yet there were many studies above this cut-off which employed plunge-freezing. The maximal tissue dimensions vitrified with plunge freezing were for human brain tissue (100 μm x 2000 μm) and *C. elegans* embryos (30 μm x 50 μm), both in the presence of cryoprotectant^22,45^. Hence, cell *size* is the fundamental driver for choice of freezing approach and appears to have greater weighting than the cellular *composition*. While this may seem obvious to experienced cryo-ET scientists, these data will help streamline new users towards viable methodological combinations for specific cell types. One noteworthy subtlety was that cryoprotectant use dominated in samples with dimensions >200 μm, effectively extending the upper limit considered viable for vitrification (Fig. 2A, black outline).

**Figure 2:**
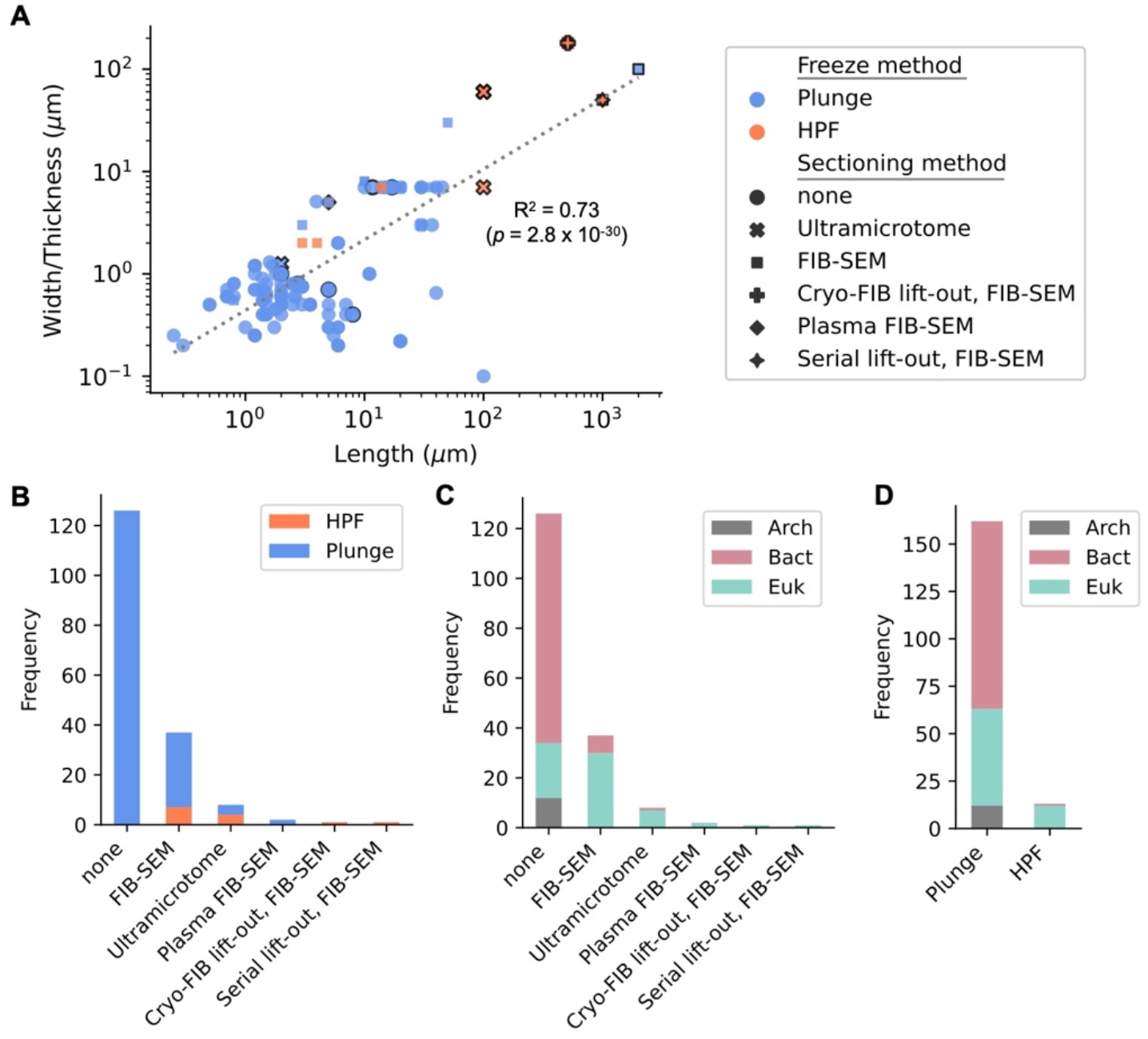
Cryo-ET methods used to freeze and section cellular material. **A)** Scatter plot of the cell/tissue length versus width/thickness for the 175 entries within this study, coloured by freezing method (plunge-freezing: blue, HPF: orange). Scatter plot marker types indicate the sectioning method used (**●**: none, x: ultramicrotome, **■**: FIB-SEM, **◆**: plasma-FIB, **✚**: cryo-FIB lift-out, FIB-SEM, **✦**: serial lift-out, FIB-SEM). A solid marker outline indicates studies in which cryoprotectant was used. **B)** Distribution of cryo-ET sectioning methods coloured by freezing type or **C)** domain of life. **D)** Distribution of cryo-ET freezing methods coloured by domain of life.

We next compared sectioning and freezing methods across studies to identify potential correlations between cell types and methods. Sectioning methods included intact cells (none), FIB-SEM, plasma-FIB, ultramicrotome, cryo-FIB lift-out, serial lift-out or a combination, while freezing methods included either plunge freezing or HPF. In most cases cells were not sectioned (72% of total, Fig. 2B), vitrified by plunge freezing, and bacterial (73% of non-sectioned entries) (Fig. 2B-C). The next most common approaches were FIB-SEM (21%) followed by the ultramicrotome (5%) and plasma FIB-SEM (1%) (Fig. 2B). Of these, eukaryotic cells dominated usage (84% of sectioned cells), and HPF was used for 19% of FIB-SEM and 50% of ultramicrotome sectioned cells (Fig. 2B-D). Plasma-FIB data is sparsely represented in this meta-analysis as it has only recently begun to be used for biological material. Finally, there were two studies which used cryo-FIB lift-out or serial lift-out followed by the FIB-SEM to study extremely large eukaryotic tissues; *D. melanogaster* eggs and *C. elegans* respectively, both of which were frozen by HPF^21,22^ (Fig. 2B-D).

### Quantification of the Fraction of Cellular Material Captured within Lamellae across the Biosphere

Cryo-ET lamellae represent only a tiny fraction of total cellular material, estimated at 0.5-4% for a typical mammalian cell^26^. To provide a more granular assessment we quantified lamellae fractions across all entries to ultimately assess how much cellular material is captured for individual cell types and organisms. We first calculated the fraction of cell/tissue material imaged assuming one lamella had been produced, with a typical lamellae width of 0.18 μm for FIB-milled samples^28^. We then divided this by the maximal length dimension from Fig. 2A. For bacterial and archaeal cells, the median percentage of cellular material imaged was 9% and 15% respectively, whereas for eukaryotes this was significantly lower (1%) (Fig. 3A). In extreme cases, only 0.009-0.6% of eukaryotic cellular material were represented in lamellae. These data align with estimates of lamellae fractions within the literature^26,46^ but are the first quantification across multiple domains of life and for individual cell types (Supplementary Table 1). This quantification shows that while lamellae from bacterial and archaeal cells capture an order of magnitude more cellular volume than those from eukaryotes, they still represent only a small fraction of the whole cell volume. This spotlights the need for complementary and/or correlated contextualising experiments^10,47^ to build statistically robust structure-function models^48^.

**Figure 3:**
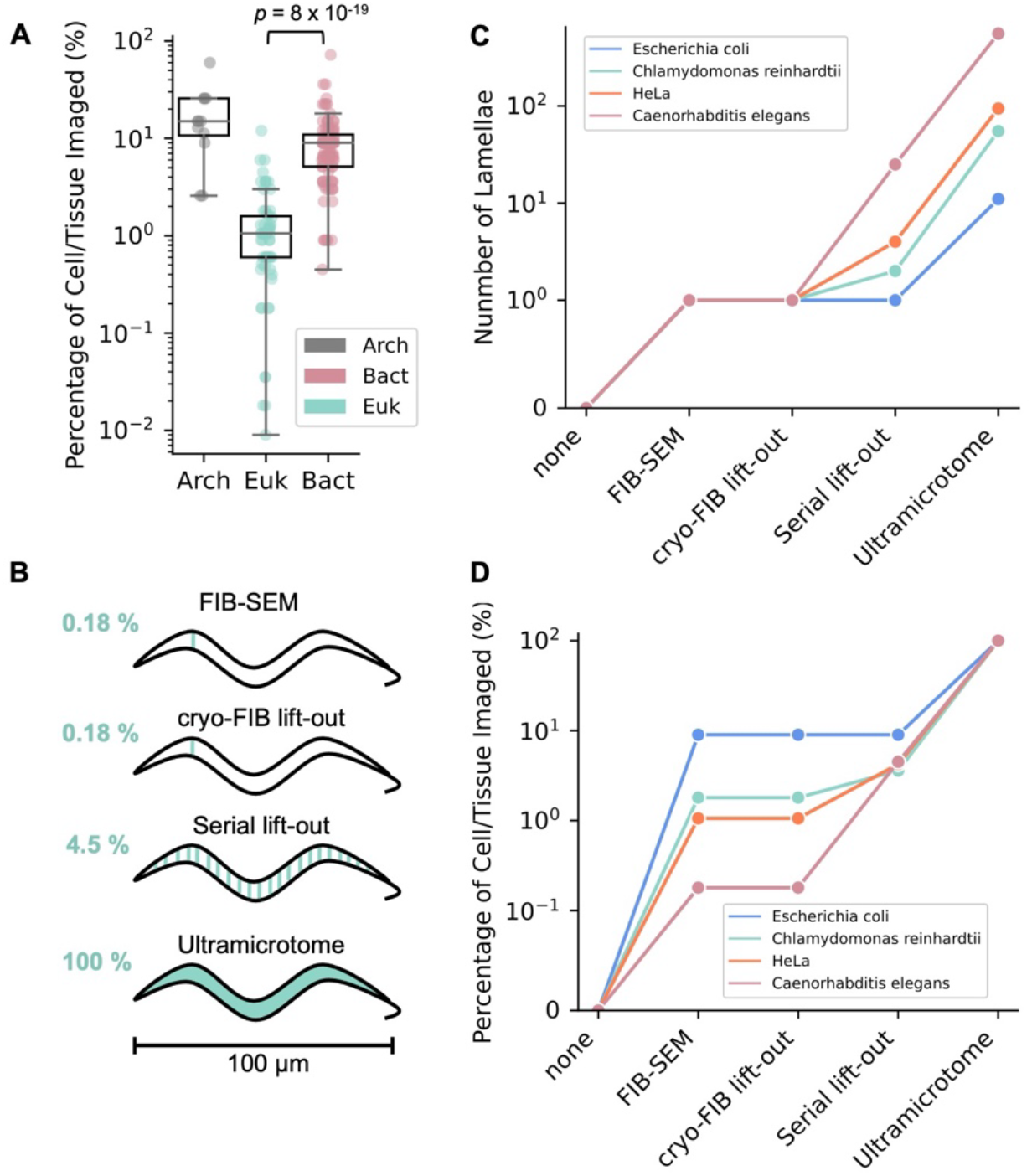
The fraction of cellular material captured within lamellae and by different sectioning methods. **A)** The percentage of cellular material captured within a lamella for each cell/tissue type across bacterial (red), eukaryotic (teal) and archaeal (grey) domains. The percentage image was defined as in Equation 1 whereby a standard lamellae width (*l* = 0.18 *μm*) was divided by the maximal dimension from Fig. 2A for each cell/tissue. Significance was tested using a Kruskal-Wallis test (*p* = 3 × 10^−21^) followed by a pairwise Dunn’s test between domains (Bact-Euk: *p* = 8 × 10^−19^, Bact.-Arch: *p* = 2 × 10^−1^, Euk-Arch: *p* = 5 × 10^−10^). **B)** Schematic of percentage of cellular material captured within lamellae by following each sectioning method applied to *C. elegens*. Teal lines indicate individual lamellae. **C)** Number of lamellae which could be produced by following each sectioning approach to completion once, for four model cells/organisms (*E. coli*: blue, *C. reinhardtii*: teal, HeLa: orange, *C. elegens*: red). **D)** Percentage of cell/tissue captured within lamellae by following each sectioning method to completion for the four cells/organisms in **C** (defined as in Equation 1).

Next, we calculated the theoretical maximum number of lamellae which could be produced for four commonly studied organisms/cells (*Escherichia coli, Chlamydomonas reinhardtii*, HeLa and *Caenorhabditis elegans*) assuming a standard workflow was followed to completion once for each of (i) no sectioning, (ii) on-grid FIB-SEM, (iii) cryo-FIB lift-out, (iv) serial lift-out and (v) the ultramicrotome (Fig. 3B). For all four cell types the cell/tissue width is too large for imaging without sectioning but one lamella is produced by following standard FIB-SEM and cryo-FIB lift-out procedures (Fig. 3C). The biggest methodological gain, both for the number of lamellae produced (Fig. 3C) and the percentage of cell/tissue material imaged (Fig. 3D) comes from switching to a serial lift-out or ultramicrotome procedure, especially for the largest organism considered here, *Caenorhabditis elegans* (Fig. 3B-D). There is really no gain from using cryo-FIB lift-out alone compared to standard FIB-SEM unless this procedure is serially coupled. The ultramicrotome can produce the highest fraction of cellular volumes by eliminating the dead space between sections. However, the diamond knife can induce artefacts during cutting, severely limiting the use of ultramicrotome sectioning for perturbation-free cryo-ET imaging^24^. For serial lift-out chunks of tissue are separated by ∼4 μm and then each are thinned to produce a lamella with pristinely preserved native architectures. Hence, we highlight serial lift-out as an emerging and powerful, approach for cryo-ET of complex tissues, especially for understanding architectural changes across whole organisms^22^.

### Extensive and Diverse Clades of Life Remain Unexplored by Cryo-ET

One goal of this analysis was to assess the explored and unexplored search space of cellular diversity in cryo-ET studies. First, to assess which cells/tissues are frequently studied via cryo-ET we calculated the number of entries for each cell/tissue type across domains of life (Fig. 4A, Supplementary Table 1). There were 55 species of bacteria represented within our analyses, of which easily culturable, genetically tractable and thin species such as *E. coli, C. crescentus* and *B. burgdorferi* were the most frequently represented (Fig. 4B). For eukaryotes, 35 species or cell lines were represented, and the most frequent occurrences were for single celled eukaryotes such as *S. cerevisiae* or widely used cell lines (HeLa, HEK293) (Fig. 4A-B). In almost all cases that did not use cell lines, studies were restricted to model organisms (*S. cerevisiae, D. melanogaster, C. elegans*). For archaea, there were 10 species represented, but studies were sparse and did not account for any of the most frequently studied model organisms (Fig. 4B).

**Figure 4:**
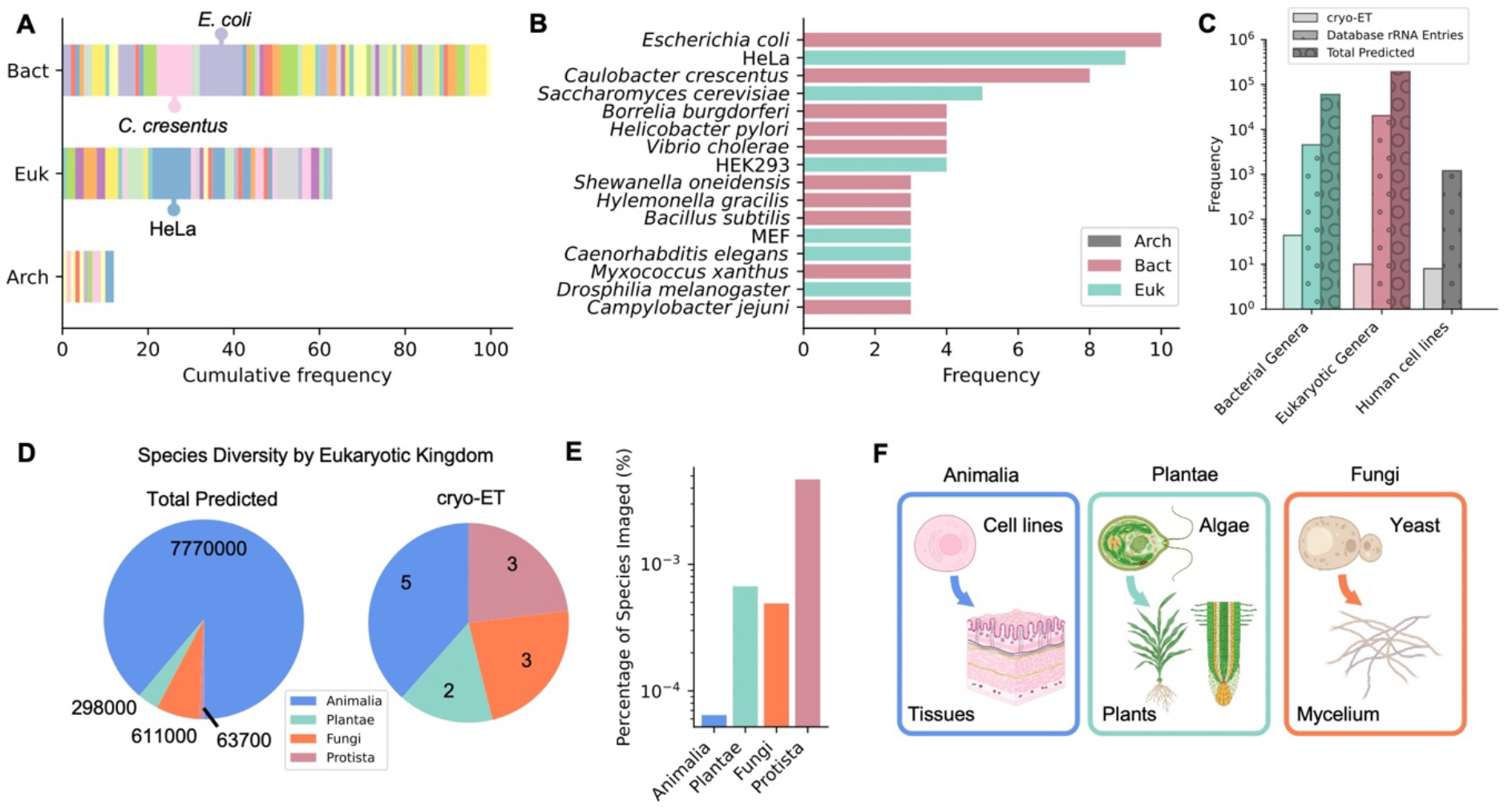
The biodiversity of cryo-ET studies compared to globally. **A)** Cumulative frequency of cell/tissue types for all 175 eukaryotic, bacterial and archaeal cryo-ET studies coloured by individual species or cell lines. Entries are in alphabetical order and correspond to those in Supplementary Table 1. The top three cell types are marked. **B)** Top species or cell lines using in cryo-ET studies. **C)** Comparison of the number of distinct bacterial genera (teal), eukaryotic genera (red) and human cell lines (grey) within cryo-ET studies (solid bars), rRNA databases (dots) and the total predicted on Earth (circles). The number of sequenced prokaryotic and eukaryotic genera were obtained from the PATRIC and EUKARYOME databases respectively^29,32^. Estimates of total richness were obtained from the SILVA and IRMNG registers^30,33^. The number of total human cell lines were derived from ^35,36^. **D)** Comparison of the number of distinct eukaryotic species predicted globally^34^ versus used in cryo-ET studies (excluding cell lines) across animalia, plantae, fungi and protista kingdoms. **E)** Percentage of distinct eukaryotic species in cryo-ET studies verses total predictions for each kingdom. **F)** Summary of under-explored eukaryotic clades of life to guide future cryo-ET studies.

Next, we compared the number of genera within our set of representative cryo-ET studies to those annotated within rRNA databases and to the total number of predicted genera on Earth (Fig. 4C). The goal of this analysis was to assess roughly how much of the biosphere is currently represented within cryo-ET studies and pinpoint unexplored areas of life. For bacteria and eukaryotes, the number of genera imaged were 2-3 orders of magnitude lower than those in sequence libraries^29,32^, and 4-5 orders of magnitude lower than the predicted planetary abundances^30,31,33^. We performed a parallel comparison for distinct human cell lines studied by cryo-ET and within rRNA databases, which also differed by 3 orders of magnitude^35,36^ (Fig. 4C). Moreover, the number of human cell lines represented was almost identical to the total diversity of eukaryotic genera studied by cryo-ET. In summary, the current biodiversity of cryo-ET studies is low and there is ample scope for expanding species representation.

To dissect whether eukaryotic studies (excluding cell lines) proportionally reflected the distribution of life on Earth, we compared the total predicted number of species in the kingdom Eukarya to the number of distinct species imaged by cryo-ET (Fig. 4D). Animalia are the most diverse eukaryotic kingdom, predicted to account for 89% of total species richness (most of which are arthropods)^34^ and yet accounted for only 38% of cryo-ET studies by kingdom. In contrast plantae, fungi and protists are predicted to account for 3%, 7% and 1% respectively^34^ but were proportionally much higher in cryo-ET studies (plantae: 15%, fungi: 23%, protista: 23%). Hence, the species richness of eukaryotic kingdoms in cryo-ET studies does not mirror the abundances of life on Earth and accounts for only a tiny proportion of the totals predicted (0.00006-0.005%) (Fig. 4E).

For animals, cryo-ET studies of human, mice and macaque tissues have been conducted in addition to the model organisms, *C. elegans* and *D. melanogaster*^45,48–50^. For fungi, yeasts have been studied in their non-filamentous forms (*S. cerevisiae*), in addition to species within the microsporidium division (*Anncaliia algerae, Encephalitozoon hellem*)^20,51,52^. Plant studies were restricted to unicellular green algae (*C. reinhardtii, Ostreococcus tauri*)^53,54^ while protists were parasites (*Plasmodium berghei, Toxoplasma gondii, Trypanosoma brucei*)^7,55–57^ or oceanic coccolithophores (*Gephyrocapsa huxleyi*)^13^. Hence, outside of the animal kingdom, eukaryotic cryo-ET studies have only been performed on unicellular organisms, illuminating a path towards future cryo-ET studies on diverse multicellular organisms which remain architecturally enigmatic.

## Discussion

### Current and Future Trajectories of Cryo-ET Methodology

Our assessment of cryo-ET data provides an integrated consortium of freezing and sectioning approaches to cellular material across all domains of life. This foundation has primed researchers to explore a wealth of cellular anatomies as the adoption of cryo-ET continues to gain momentum (Fig. 1). Hereafter, we outline a series of methodological principles to guide future cryo-ET studies and accelerate *in situ* cellular discovery.

We aggregated the freezing and sectioning approaches used for cells/tissue across all domains of life (Fig. 2). These data are intended to benefit the community by reducing the time and cost associated with optimisation of samples, enhance the accessibility of cryo-ET for new users and pinpoint appropriate pairwise combinations of vitrification and sectioning methods. We also provide a comprehensive breakdown of preparation conditions for individual cells/tissues to compare or expand upon established studies (Supplementary Table 1). Our analyses show that bacterial and archaeal cells can be routinely plunge-frozen and require minimal to no sectioning, provided cells are sparsely distributed on the grid and are hydrated (Fig. 2A-D). For bacterial cells thicker than 300 nm, FIB-SEM sectioning is the primary methodological approach. Increased cell density or thickness of the hydrated layer, however, may be required to prevent grid buckling during milling. For unicellular eukaryotic cells, the most frequent pipeline was plunge-freezing followed by gallium or plasma FIB milling (Fig. 2A-D). Once cell/tissue dimensions surpass ∼10-100 μm it is usually appropriate to switch from plunge-freezing to HPF and the vitrification threshold can be extended beyond 200 μm by cryoprotectant use (Fig. 2A).

Expanding upon these foundational principles, we highlight several powerful approaches which push the boundaries of current cryo-ET methodology. Firstly, plasma-FIB use was underrepresented in our analysis because the methodology is still emerging. We anticipate plasma-FIB uptake to increase, especially for eukaryotic cells, as material can be sectioned quickly and may have reduced sample damage compared to the traditional gallium FIB^19^. Secondly, a specialised use of HPF is to freeze *bacterial* layers via the ‘Waffle method’^51^. This would be highly valuable for studying multiple cells within a single lamella *en mass* and understanding how cells are situated within their surrounding environment e.g. in biofilms where cell-cell and cell-extracellular matrix interactions are critical^58^. Next, our assessment of the potential for different sectioning approaches to image greater proportions of cells than the 1-10% typically imaged (Fig. 3) identified serial lift-out as a high-potential method^21–23^. This approach had the greatest fractional change for larger, eukaryotic cells/tissues compared to FIB-SEM and standard cryo-FIB lift-out approaches but was only represented once within our meta-analyses^22^. Since 2024, serial lift-out has been successfully used to obtain vitrified plant protonemata from *Physcomitrium patens* with an adapted single-sided needle attachment for faster manipulation^46^. These studies are a substantial advancement for studying (i) plant tissues and (ii) all multicellular life because they highlight the increased biological insight that is obtained by tracing structural architectures along the same organism. For example, serial lift-out applied to *C. elegans* revealed new packing arrangements of actin-myosin bundles along larvae which could not be inferred from individual lamellae^22^. We are on the precipice of being able to visualise structures across length-scales to trace bio-anatomical features across whole organisms. For processes where chemical and environmental gradients are important for structural emergence, such as in developmental biology, molecular scale resolution across these axes will be powerful.

### Chartering Underexplored Lineages in Structural Cell Biology

After identifying methodology to better capture structural landscapes, the question remains which organisms to target for maximising biological discovery. Across kingdoms, the biodiversity captured by cryo-ET is 2-5 orders of magnitude lower than those in sequence databases or on Earth (Fig. 4). Understandably, most cryo-ET studies have been performed on unicellular life or cell lines due to requirements for thin samples and ease of culturing (Fig. 4A-B). While commonly studied organisms are useful for method development and benchmarking^3,47,49^, we must be cautious not to create too much redundancy or repetition bias within the literature (Fig. 4B). For example, studying how architectures differ between species can inform functional and evolutionary trends. This is evidenced by recent cryo-ET structures of pyrenoid compartments from the diatoms *Phaeodactylum tricornutum* and *Thalassiosira psuedonana* which have distinct thylakoid sheets and rubisco partitioning that may alter gaseous diffusion compared to in well studied *C. reinhardtii* pyrenoids^59^. Moreover, these data posit an urgent need to assess whether the immortalized nature of eukaryotic cell lines results in architectures which may not be reflective of healthy cells or tissues^60^. Hence, expanding beyond model organisms is functionally necessary for revealing untapped cellular information by cryo-ET (Fig. 4C).

We identify unexplored clades of life where future cryo-ET studies would be welcomed (Fig. 4F). For eukaryotes, animalia tissues, embryophyte plantae and filamentous fungi are ripe for exploration. Recent, pioneering studies of human and mouse brain tissue revealed the arrangement of fibrilla plaques implicated in Alzheimers^45,50^ and the *in situ* location of synaptic glutamate receptors^61^. Importantly, these explorations of animal tissues were only made possible by advances in sample manipulation with cryogenic needles, as discussed above. One critical dimension that remains underexplored is imaging multi-species interactions in disease^56^ and ecological contexts^62^. For example, a study of the *Plasmodium falciparum* infection cycle in human erythrocytes showed altered ribosomal abundances when parasites were first treated with drugs^63^. This highlights how cryo-ET can be used to assess how multicellular landscape are shifted by pharmaceutical or environmental intervention. Hence, we are poised to enter an era of cryo-ET discovery of cellular interactions in real space however advances in culturing methods will be necessary. For example, many multi-species communities important to ecological functioning (such a lichens and seaweeds) are not easily cultured in labs or in liquid media compatible with freezing. Concentrated salt water has a depressed freezing point which hinders sample vitrification^64^ and methods do not currently exist for growing mineral associated communities on grids. Development of novel culturing methods, paired with lift-out techniques and customized grids, are crucial to accelerate cryo-ET discovery across biodiverse lineages.

Finally, coupling cryo-ET data with additional contextualising modality will increase the functional impact of structural cell biology^65,66^. Molecular context can be gained through conjugated gold^61,67^, cathodoluminescent labels^68^ or genetically encoded fluorophores to pinpoint individual molecules *in situ*^10,69^. Additionally, micro-fluidics-based mix-and-spray approaches allow capture of cellular processes with temporal context^47^. This approach was, for example, used to track pH induced shedding of the *C. crescentus* S-layer^47^. These bonus methods add colour to cryo-ET data, enabling more robust structure-function relationships to emerge than in traditional, label-free tomograms.

Herein, we outline the most comprehensive synthesis of cellular cryo-ET data to date. We identify critical transition points between freezing and sectioning approaches and the underlying factors which shape methodological choice. We provide an exhaustive comparison of the cellular fractions within lamellae across all domains of life and quantitatively evaluate how close current sectioning methods get to imaging whole cells or organisms. We establish a series of guiding principles for cellular cryo-ET research to reduce times and cost associated with imaging, improve cryo-ET accessibility and facilitate constructions of structural architectures over multicellular communities. Finally, we identify vast clades of life where structural information is lacking and where cryo-ET research can expedite our understanding of cellular behaviour in space.

## Acknowledgements

T.B.A. was supported by Schmidt Science Fellows in partnership with the Rhodes Trust. L.B. was supported by a Stanford Doerr School of Sustainability Discovery grant awarded to L.B. and K.G.P. K.G.P was supported by US Department of Energy grant DE-SC0023661 and is a Fellow of the Canadian Institute for Advanced Research Fungal Kingdom program. P.D.D was supported by US Department of Energy grant DE-AC02-76SF00515.

## Author contributions

T.B.A and L.B. conceptualised the study. T.B.A. conducted analyses and wrote the paper with input from all authors.

