## Supplementary Information for "Cryogenic Electron Tomography by the Numbers: Charting Underexplored Lineages in Structural Cell Biology"

T. Bertie Ansell[1,2,], Louis Berrios[1], Kabir Peay[1,3], Peter Dahlberg[2]
[1] Department of Biology, Stanford University, Stanford, CA 94305, USA
[2] Division of Cryo-EM and Bioimaging, SLAC National Accelerator Laboratory, Menlo Park, CA 94025, USA
[3] Department of Earth System Science, Stanford University, Stanford, CA 94305, USA

### Supplementary Table 1: Cells and tissues studied by cryo-ET included within this analysis.

| Cell type | Domain | Freezing method | Sectioning method | Cryoprotectant | Cell Width/ Thickness (µm) | Cell Length (µm) | Cell Length-2 (µm) | Cell Dimensions PMID/DOI | Percentage of Cell/Tissue Imaged (%) | Shape | Reference |
|---|---|---|---|---|---|---|---|---|---|---|---|
| Aplasma | Arch | Plunge | none | | 0.6 | 0.7 | | 23865623 | 25.71 | sphere | 1 |
| ARMAN | Arch | Plunge | none | | 0.2 | 0.3 | | 18946497 | 60.00 | sphere | 2 |
| Eplasma | Arch | Plunge | none | | 0.6 | 0.7 | | 23865623 | 25.71 | sphere | 1 |
| Ferroplasma | Arch | Plunge | none | | 0.6 | 0.7 | | 23865623 | 25.71 | sphere | 1 |
| Halobacterium salinarum | Arch | Plunge | none | | 0.9 | 1.4 | | 34594449 | 12.86 | capsule | 3 |
| Haloquadratum walsbyi | Arch | Plunge | none | | 1.2 | 1.2 | 0.2 | 34594449 | 15.00 | rectangle | 4 |
| Methanoregula formicica | Arch | Plunge | none | | 0.4 | 2 | | 34594449 | 9.00 | capsule | 3 |
| Methanospirillum hungatei | Arch | Plunge | none | | 0.5 | 7 | | 39671499 | 2.57 | capsule | 5 |
| Methanosprillum hungatei | Arch | Plunge | none | | 0.4 | 7 | 0.4 | 39671499 | 2.57 | rectangle | 3 |
| Sulfolobus acidocaldarius | Arch | Plunge | none | | 1.3 | 1.6 | | 34594449 | 11.25 | capsule | 6 |
| Thermococcus kodakarensis | Arch | Plunge | none | | 1.2 | 1.2 | | 34594449 | 15.00 | sphere | 3 |
| Thermococcus kodakarensis | Arch | Plunge | none | | 1.2 | 1.2 | | 34594449 | 15.00 | sphere | 7 |
| Acetonema longum | Bact | Plunge | none | | 0.3 | 6 | | 34594449 | 3.00 | capsule | 8 |

| | | | | | | | | | | | |
|---|---|---|---|---|---|---|---|---|---|---|---|
| Acetonema longum (APO-1) | Bact | Plunge | none | | 0.3 | 6 | | 34594449 | 3.00 | capsule | [9] |
| Acinetobacter baumannii (RUH 30233T) | Bact | Plunge | Ultramicrotome | 20% glycerol | 1.25 | 2 | | 29686523 | 9.00 | capsule | [10] |
| Agrobacterium tumetaciens | Bact | Plunge | none | | 0.8 | 1.5 | | 34594449 | 12.00 | capsule | [11] |
| Amoebophilus asiaticus | Bact | Plunge | FIB-SEM | | 0.55 | 0.8 | | 34594449 | 22.50 | capsule | [12] |
| Azospirillum brasilense | Bact | Plunge | none | | 0.7 | 2 | | doi.org/10.1007/s00374-024-01839-4 | 9.00 | capsule | [13] |
| Bacillus cereus | Bact | Plunge | none | | 0.75 | 3 | | 34594449 | 6.00 | capsule | [9] |
| Bacillus subtilis | Bact | Plunge | none | | 0.75 | 3 | | 34594449 | 6.00 | capsule | [14] |
| Bacillus subtilis | Bact | Plunge | none | | 0.75 | 3 | | 34594449 | 6.00 | capsule | [9] |
| Bacillus subtilis | Bact | Plunge | none | | 0.75 | 3 | | 34594449 | 6.00 | capsule | [8] |
| Bacillus thuringiensis | Bact | Plunge | none | | 0.75 | 3 | | 34594449 | 6.00 | capsule | [9] |
| Bdellovibrio bacteriovorus | Bact | Plunge | none | | 0.4 | 1.4 | | 34594449 | 12.86 | capsule | [13] |
| Bdellovibrio bacteriovorus | Bact | Plunge | none | | 0.4 | 1.4 | | 34594449 | 12.86 | capsule | [15] |
| Borrelia burgdorferi | Bact | Plunge | none | | 0.22 | 20 | | 7043737 | 0.90 | capsule | [16] |
| Borrelia burgdorferi | Bact | Plunge | none | | 0.22 | 20 | | 7043737 | 0.90 | capsule | [17] |
| Borrelia burgdorferi | Bact | Plunge | none | | 0.22 | 20 | | 7043737 | 0.90 | capsule | [13] |
| Borrelia burgdorferi | Bact | Plunge | none | | 0.22 | 20 | | 7043737 | 0.90 | capsule | [18] |
| Borrelia garinii | Bact | Plunge | none | | 0.22 | 20 | | 7043737 | 0.90 | capsule | [16] |
| Brucella abortus | Bact | Plunge | none | | 0.7 | 1.4 | | 34594449 | 12.86 | capsule | [13] |
| Campylobacter jejuni | Bact | Plunge | none | | 0.25 | 1.2 | | 34594449 | 15.00 | capsule | [13] |
| Campylobacter jejuni | Bact | Plunge | none | | 0.25 | 1.2 | | 34594449 | 15.00 | capsule | [19] |
| Campylobacter jejuni | Bact | Plunge | none | | 0.25 | 1.2 | | 34594449 | 15.00 | capsule | [8] |
| Caulobacter crescentus | Bact | Plunge | none | 30% ethylene glycol | 0.8 | 2.75 | | 25778096 | 6.55 | capsule | [20] |
| Caulobacter crescentus | Bact | Plunge | none | | 0.8 | 2.75 | | 25778096 | 6.55 | capsule | [21] |
| Caulobacter crescentus | Bact | Plunge | none | | 0.8 | 2.75 | | 25778096 | 6.55 | capsule | [22] |
| Caulobacter crescentus | Bact | Plunge | none | | 0.8 | 2.75 | | 25778096 | 6.55 | capsule | [23] |
| Caulobacter crescentus | Bact | Plunge | none | | 0.8 | 2.75 | | 25778096 | 6.55 | capsule | [24] |
| Caulobacter crescentus | Bact | Plunge | none | | 0.8 | 2.75 | | 25778096 | 6.55 | capsule | [25] |

| | | | | | | | | | | | |
|---|---|---|---|---|---|---|---|---|---|---|---|
| Caulobacter crescentus | Bact | Plunge | none | | 0.8 | 2.75 | | 25778096 | 6.55 | capsule | 13 |
| Caulobacter crescentus | Bact | Plunge | none | | 0.8 | 2.75 | | 25778096 | 6.55 | capsule | 18 |
| Chitinophaga pinensis | Bact | Plunge | none | | 0.65 | 40 | | 21304681 | 0.45 | capsule | 18 |
| Delftia acidovorans | Bact | Plunge | none | | 0.8 | 0.8 | | 34594449 | 22.50 | sphere | 18 |
| Escherichia coli | Bact | Plunge | FIB-SEM | | 1 | 2 | | 2403552, 24287933 | 9.00 | capsule | 26 |
| Escherichia coli | Bact | HPF | FIB-SEM | | 1 | 2 | | 2403552, 24287933 | 9.00 | capsule | 27 |
| Escherichia coli | Bact | Plunge | none | | 1 | 2 | | 2403552, 24287933 | 9.00 | capsule | 13 |
| Escherichia coli | Bact | Plunge | none | 10% Ficoll PM 70 | 1 | 2 | | 2403552, 24287933 | 9.00 | capsule | 28 |
| Escherichia coli | Bact | Plunge | none | | 1 | 2 | | 2403552, 24287933 | 9.00 | capsule | 8 |
| Escherichia coli (1094) | Bact | Plunge | FIB-SEM | | 1 | 2 | | 2403552, 24287933 | 9.00 | capsule | 11 |
| Escherichia coli (K12) | Bact | Plunge | FIB-SEM | | 1 | 2 | | 2403552, 24287933 | 9.00 | capsule | 29 |
| Escherichia coli (MG1655, XL-10) | Bact | Plunge | none | | 1 | 2 | | 2403552, 24287933 | 9.00 | capsule | 25 |
| Escherichia coli (MG1655) | Bact | Plunge | FIB-SEM | | 1 | 2 | | 2403552, 24287933 | 9.00 | capsule | 30 |
| Escherichia coli (mini cells) | Bact | Plunge | none | | 0.25 | 0.25 | | 2838458 | 72.00 | sphere | 31 |
| Flavobacterium anhuiense | Bact | Plunge | none | | 0.3 | 1.75 | | 18398165 | 10.29 | capsule | 18 |
| Flavobacterium johnsoniae | Bact | Plunge | none | | 0.5 | 3 | | 33441737 | 6.00 | capsule | 18 |
| Geobacter sulfurreducens | Bact | Plunge | none | | 0.5 | 2 | | 10.1080/01490451.2015.1099765 | 9.00 | capsule | 32 |
| Gluconacetobacter hansenii | Bact | Plunge | FIB-SEM | | 0.6 | 2 | | 10.1007/s13213-011-0288-4 | 9.00 | capsule | 33 |
| Gluconacetobacter xylinus | Bact | Plunge | none | | 0.6 | 2 | | 10.1007/s13213-011-0288-4 | 9.00 | capsule | 33 |
| Halothiobacillus neapolitanus | Bact | Plunge | none | | 0.45 | 1.8 | | 34594449 | 10.00 | capsule | 34 |
| Halothiobacillus neapolitanus | Bact | Plunge | none | | 0.45 | 1.8 | | 34594449 | 10.00 | capsule | 13 |
| Helicobacter hepaticus | Bact | Plunge | none | | 0.3 | 5 | | 34594449 | 3.60 | capsule | 18 |
| Helicobacter hepaticus | Bact | Plunge | none | | 0.3 | 5 | | 34594449 | 3.60 | capsule | 8 |
| Helicobacter pylori | Bact | Plunge | none | | 0.5 | 3.5 | | 8903168 | 5.14 | capsule | 35 |
| Helicobacter pylori | Bact | Plunge | none | | 0.5 | 3.5 | | 8903168 | 5.14 | capsule | 36 |
| Helicobacter pylori | Bact | Plunge | none | | 0.5 | 3.5 | | 8903168 | 5.14 | capsule | 13 |
| Helicobacter pylori | Bact | Plunge | none | | 0.5 | 3.5 | | 8903168 | 5.14 | capsule | 18 |

| | | | | | | | | | | | |
|---|---|---|---|---|---|---|---|---|---|---|---|
| Hydrogenovibrio crunogenus | Bact | Plunge | none | | 0.5 | 1.5 | | 34594449 | 12.00 | capsule | 8 |
| Hylemonella gracilis | Bact | Plunge | none | | 0.2 | 6 | | 34594449 | 3.00 | capsule | 13 |
| Hylemonella gracilis | Bact | Plunge | none | | 0.2 | 6 | | 34594449 | 3.00 | capsule | 18 |
| Hylemonella gracilis | Bact | Plunge | none | | 0.2 | 6 | | 34594449 | 3.00 | capsule | 8 |
| Hyphomonas neptunium | Bact | Plunge | none | | 1 | 1.8 | | 34594449 | 10.00 | capsule | 13 |
| Legionella pneumophila | Bact | Plunge | none | | 0.5 | 2 | | 34594449 | 9.00 | capsule | 37 |
| Legionella pneumophila | Bact | Plunge | none | | 0.5 | 2 | | 34594449 | 9.00 | capsule | 38 |
| Magnetospirillum magnetotacticum | Bact | Plunge | none | | 0.6 | 2.6 | | 34594449 | 6.92 | capsule | 18 |
| Magnetospirillum magnetotacticum (AMB-1) | Bact | Plunge | none | | 0.6 | 2.6 | | 34594449 | 6.92 | capsule | 39 |
| Methylomicrobium alcaliphilum | Bact | Plunge | none | | 1 | 1.2 | | 34594449 | 15.00 | capsule | 40 |
| Methyloprofundus sedimenti | Bact | Plunge | none | | 1.2 | 1.7 | | 34594449 | 10.59 | capsule | 41 |
| Mycobacterium smegmatis | Bact | Plunge | none | | 0.5 | 2.5 | | 34594449 | 7.20 | capsule | 13 |
| Mycoplasma pneumoniae | Bact | Plunge | none | | 0.3 | 1 | | 34594449 | 18.00 | capsule | 42 |
| Myxococcus xanthus | Bact | Plunge | none | 5% Ficoll PM 70, 10% Ficoll PM 70 and 10% ethylene glycol | 0.7 | 5 | | 34594449 | 3.60 | capsule | 43 |
| Myxococcus xanthus | Bact | Plunge | none | | 0.7 | 5 | | 34594449 | 3.60 | capsule | 13 |
| Myxococcus xanthus | Bact | Plunge | none | | 0.7 | 5 | | 34594449 | 3.60 | capsule | 18 |
| Parachlamydia acanthamoebae | Bact | Plunge | none | | 0.5 | 0.5 | | 8979345 | 36.00 | sphere | 44 |
| Prosthecobacter debontii | Bact | Plunge | none | | 0.4 | 5 | | 9296261 | 3.60 | capsule | 13 |
| Prosthecobacter fluviatilus | Bact | Plunge | none | | 0.5 | 5 | | 18599695 | 3.60 | capsule | 13 |
| Prosthecobacter vanneervenii | Bact | Plunge | none | 10% Ficoll PM 70 | 0.4 | 8 | | 9296261 | 2.25 | capsule | 28 |
| Prosthecobacter vanneervenii | Bact | Plunge | none | | 0.4 | 8 | | 9296261 | 2.25 | capsule | 13 |
| Proteus mirabilis | Bact | Plunge | none | | 1 | 2 | | 15213138 | 9.00 | capsule | 45 |
| Protochlamydia amoebophila | Bact | Plunge | none | | 0.8 | 0.8 | | 24292151 | 22.50 | sphere | 44 |
| Pseudoalteromonas luteoviolacea | Bact | Plunge | none | | 0.6 | 1.5 | | 34594449 | 12.00 | capsule | 18 |
| Pseudomonas aeruginosa | Bact | Plunge | none | | 0.8 | 2 | | 34594449 | 9.00 | capsule | 40 |
| Ralstonia eutropha | Bact | Plunge | none | | 0.7 | 1.2 | | 22178974 | 15.00 | capsule | 13 |

| | | | | | | | | | | | |
|---|---|---|---|---|---|---|---|---|---|---|---|
| Ralstonia eutropha (H16) | Bact | Plunge | none | | 0.7 | 1.2 | | 22178974 | 15.00 | capsule | [46] |
| Rectangular bacterial cells | Bact | Plunge | none | | 5.08 | 3.95 | | 37055390 | 4.56 | rectangle | [47] |
| Salmonella enterica (mini cells) | Bact | Plunge | none | | 0.5 | 0.5 | | 24284544 | 36.00 | sphere | [17] |
| Shewanella oneidensis | Bact | Plunge | none | | 0.6 | 2 | | 34594449 | 9.00 | capsule | [48] |
| Shewanella oneidensis | Bact | Plunge | none | | 0.6 | 2 | | 34594449 | 9.00 | capsule | [40] |
| Shewanella oneidensis | Bact | Plunge | none | | 0.6 | 2 | | 34594449 | 9.00 | capsule | [18] |
| Shewanella putrifaciens | Bact | Plunge | none | | 0.6 | 2 | | 34594449 | 9.00 | capsule | [13] |
| Simkania negevensis | Bact | Plunge | none | | 0.7 | 0.7 | | 34594449 | 25.71 | sphere | [44] |
| Tetrasphaera remsis | Bact | Plunge | none | | 0.6 | 0.8 | | 34594449 | 22.50 | capsule | [49] |
| Thiomicrospira crunogena | Bact | Plunge | none | | 0.4 | 1.5 | | 10.1099/00207713-35-4-422 | 12.00 | capsule | [34] |
| Thiomicrospira crunogena | Bact | Plunge | none | | 0.4 | 1.5 | | 10.1099/00207713-35-4-422 | 12.00 | capsule | [13] |
| Thiomonas intermedia | Bact | Plunge | none | | 0.55 | 1.4 | | 34594449 | 12.86 | capsule | [34] |
| Thiomonas intermedia | Bact | Plunge | none | | 0.55 | 1.4 | | 34594449 | 12.86 | capsule | [13] |
| Treponema primitia | Bact | Plunge | none | | 0.25 | 5.5 | | 34594449 | 3.27 | capsule | [50] |
| Vibrio cholerae | Bact | Plunge | none | | 0.8 | 2.5 | | 26933214 | 7.20 | capsule | [51] |
| Vibrio cholerae | Bact | Plunge | none | | 0.8 | 2.5 | | 26933214 | 7.20 | capsule | [52] |
| Vibrio cholerae | Bact | Plunge | none | | 0.8 | 2.5 | | 26933214 | 7.20 | capsule | [13] |
| Vibrio cholerae | Bact | Plunge | none | | 0.8 | 2.5 | | 26933214 | 7.20 | capsule | [40] |
| Yersinia enterocolitica | Bact | Plunge | none | | 0.65 | 2 | | 22019131 | 9.00 | capsule | [21] |
| 17Cl-1 (mouse) | Euk | Plunge | FIB-SEM | | 7 | 35 | | 31536774 | 0.51 | cone | [53] |
| Anncaliia algerae | Euk | HPF | FIB-SEM | | 2 | 3 | | 31332877 | 6.00 | capsule | [27] |
| Anncaliia algerae (microsporidia tubule region) | Euk | Plunge | none | | 0.1 | 100 | | 31332877 | 0.18 | capsule | [54] |
| BSC-1 (monkey kidney) | Euk | Plunge | FIB-SEM | | 7 | 30 | | 11222860 | 0.60 | cone | [55] |
| BSC-1 (monkey kidney) | Euk | Plunge | none | | 7 | 30 | | 11222860 | 0.60 | cone | [56] |
| Caenorhabditis elegans | Euk | HPF | FIB-SEM | 2-methyl pentane | 50 | 1000 | | 25961413 | 0.02 | capsule | [57] |
| Caenorhabditis elegans | Euk | HPF | Serial lift-out, FIB-SEM | 20% Ficoll 400 | 50 | 1000 | | 25961413 | 0.02 | capsule | [58] |

| | | | | | | | | | | | |
|---|---|---|---|---|---|---|---|---|---|---|---|
| Caenorhabditis elegans (embrios) | Euk | Plunge | FIB-SEM | | 30 | 50 | | 25961413 | 0.36 | sphere | [57] |
| Chlamydomonas reinhardtii | Euk | Plunge | FIB-SEM | | 8 | 10 | | 11337403 | 1.80 | capsule | [59] |
| Chlamydomonas reinhardtii | Euk | Plunge | FIB-SEM | | 8 | 10 | | 11337403 | 1.80 | capsule | [60] |
| Drosphilia melanogaster (eggs) | Euk | HPF | Cryo-FIB lift-out, FIB-SEM | 20% Ficoll 70 | 180 | 510 | | 19032497 | 0.04 | sphere | [60] |
| Drosphilia melanogaster (embrios) | Euk | HPF | FIB-SEM | 2-methyl pentane | 180 | 510 | | 19032497 | 0.04 | sphere | [57] |
| Drosphilia melanogaster (S2 cells) | Euk | Plunge | none | 0.125% DMSO | 7 | 11.7 | | 29907103 | 1.54 | cone | [61] |
| Encephalitozoon hellem | Euk | HPF | FIB-SEM | | 2 | 4 | | 35387991 | 4.50 | sphere | [27] |
| Gephyrocapsa huxleyi | Euk | Plunge | FIB-SEM | | 3 | 3 | | 31530807 | 6.00 | sphere | [60] |
| HEK293 | Euk | HPF | Ultramicrotome | | 7 | 14 | | https://bionumbers.hms.harvard.edu/files/Sizes%20of%20various%20cells.pdf | 1.29 | cone | [62] |
| HEK293 | Euk | Plunge | FIB-SEM | | 7 | 14 | | https://bionumbers.hms.harvard.edu/files/Sizes%20of%20various%20cells.pdf | 1.29 | cone | [55] |
| HEK293 (HEK293S) | Euk | HPF | FIB-SEM | | 7 | 14 | | https://bionumbers.hms.harvard.edu/files/Sizes%20of%20various%20cells.pdf | 1.29 | cone | [27] |
| HEK293 (HEK293T) | Euk | Plunge | none | | 7 | 14 | | https://bionumbers.hms.harvard.edu/files/Sizes%20of%20various%20cells.pdf | 1.29 | cone | [63] |
| HeLa (CCL-2) (human cervical cancer) | Euk | Plunge | Plasma FIB-SEM | glycerol | 7 | 17 | | 5761872 | 1.06 | cone | [64] |
| HeLa (human cervical cancer) | Euk | Plunge | FIB-SEM | | 7 | 17 | | 5761872 | 1.06 | cone | [29] |
| HeLa (human cervical cancer) | Euk | Plunge | FIB-SEM | | 7 | 17 | | 5761872 | 1.06 | cone | [60] |
| HeLa (human cervical cancer) | Euk | Plunge | none | EAFS (ethylene glycol, acetamide, Ficoll, sucrose), DES (DMSO, ethylene glycol, sucrose, FCS), DE (same minus sucrose), dextran | 7 | 17 | | 5761872 | 1.06 | cone | [65] |
| HeLa (human cervical cancer) | Euk | Plunge | none | | 7 | 17 | | 5761872 | 1.06 | cone | [66] |
| HeLa (human cervical cancer) | Euk | Plunge | none | | 7 | 17 | | 5761872 | 1.06 | cone | [67] |
| HeLa (human cervical cancer) | Euk | Plunge | none | | 7 | 17 | | 5761872 | 1.06 | cone | [68] |
| HeLa (human cervical cancer) | Euk | Plunge | FIB-SEM | | 7 | 17 | | 5761872 | 1.06 | cone | [69] |

| | | | | | | | | | | | |
|---|---|---|---|---|---|---|---|---|---|---|---|
| HeLa (human cervical cancer) | Euk | Plunge | FIB-SEM | | 7 | 17 | | 5761872 | 1.06 | cone | 70 |
| HepG2 (human liver cancer) | Euk | Plunge | FIB-SEM | | 7 | 15.5 | | 34884942 | 1.16 | cone | 69 |
| HL-1 (mouse cardiomyocyte) | Euk | HPF | Ultramicrotome | 20% dextran | 7 | 100 | 20 | 9501201 | 0.18 | rectangular based pyramid | 71 |
| Human brain tissue | Euk | Plunge | FIB-SEM | 20% glycerol, 1M trehalose in DPBS | 100 | 2000 | 2000 | 38531877 | 0.01 | rectangle | 72 |
| Human primary adiposite cells | Euk | Plunge | none | | 7 | 20 | | 29641234 | 0.90 | cone | 66 |
| HUVEC (human umbilical vein endothelial cells) | Euk | Plunge | none | | 7 | 17 | | https://www.thermofisher.com/content/dam/LifeTech/migration/en/filelibrary/cell-tissue-analysis/pdfs.par.12078.file.dat/huvec.pdf | 1.06 | cone | 73 |
| INS-1E (rat insulinoma beta cells) | Euk | Plunge | none | | 7 | 45 | | 29078993 | 0.40 | cone | 66 |
| iPSC neurons | Euk | Plunge | none | | 7 | 15 | | 20807017 | 1.20 | cone | 74 |
| MDCK (canine kidney) | Euk | Plunge | FIB-SEM | | 7 | 30 | | 25606673 | 0.60 | cone | 55 |
| MEF (MEF-IRE1a-mNG) (mouse embryonic fibroblasts) | Euk | Plunge | none | | 3 | 30 | 10 | 34088673 | 0.60 | rectangular based pyramid | 75 |
| MEF (mouse embryonic fibroblasts) | Euk | Plunge | Ultramicrotome | | 3 | 30 | 10 | 34088673 | 0.60 | rectangular based pyramid | 10 |
| MEF (mouse embryonic fibroblasts) | Euk | Plunge | FIB-SEM | | 3 | 30 | 10 | 34088673 | 0.60 | rectangular based pyramid | 76 |
| Mouse brain tissue | Euk | HPF | Ultramicrotome | 20% w/v 40000 Dextran in NMDG-HEPES solution | 60 | 100 | 150 | 37198197 | 0.18 | rectangle | 77 |
| Mouse brain tissue (2mm) | Euk | HPF | Ultramicrotome | 20% w/v 40000 Dextran in NMDG-HEPES solution | 60 | 100 | 150 | 37198197 | 0.18 | rectangle | 78 |
| NIH/3T3 (mouse embryonic fibroblast) | Euk | Plunge | FIB-SEM | | 7 | 15 | | https://bionumbers.hms.harvard.edu/files/Sizes%20of%20various%20cells.pdf | 1.20 | cone | 55 |
| Ostreococcus tauri | Euk | Plunge | none | | 0.65 | 1.5 | | 17710148 | 12.00 | capsule | 79 |
| P19 (mouse embryonic cancer) | Euk | Plunge | Ultramicrotome | | 7 | 20 | | 20163207 | 0.90 | cone | 71 |
| P19 (mouse embryonic cancer) | Euk | Plunge | FIB-SEM | | 7 | 20 | | 20163207 | 0.90 | cone | 59 |
| PC3 (human prostate cancer) | Euk | Plunge | FIB-SEM | | 7 | 20 | | 26241348 | 0.90 | cone | 80 |
| Plasmodium berghei (sporozite) | Euk | Plunge | none | | 1 | 11 | | 28108531 | 1.64 | capsule | 81 |
| Plasmodium berghei (sporozite) | Euk | Plunge | none | | 1 | 11 | | 28108531 | 1.64 | capsule | 82 |
| PtK-1 (rat kangaroo epithelial kidney) | Euk | Plunge | FIB-SEM | | 7 | 30 | | 36539423 | 0.60 | cone | 83 |

| | | | | | | | | | | | |
|---|---|---|---|---|---|---|---|---|---|---|---|
| PtK-2 (potoroo epithelial kidney) | Euk | Plunge | none | | 3 | 37 | | 39308425 | 0.49 | cone | [84] |
| Rhesus macaque fibroblasts | Euk | Plunge | none | | 7 | 40 | | 29078993 | 0.45 | cone | [66] |
| Saccharomyces cerevisiae | Euk | Plunge | FIB-SEM | | 5 | 5 | | 36707648 | 3.60 | sphere | [60] |
| Saccharomyces cerevisiae | Euk | Plunge | FIB-SEM | | 5 | 5 | | 36707648 | 3.60 | sphere | [85] |
| Saccharomyces cerevisiae | Euk | Plunge | Plasma FIB-SEM | glycerol | 5 | 5 | | 36707648 | 3.60 | sphere | [64] |
| Saccharomyces cerevisiae | Euk | HPF | FIB-SEM | | 5 | 5 | | 36707648 | 3.60 | sphere | [27] |
| Saccharomyces cerevisiae | Euk | Plunge | FIB-SEM | | 5 | 5 | | 36707648 | 3.60 | sphere | [56] |
| Sum159 (human breast cancer) | Euk | Plunge | FIB-SEM | | 7 | 40 | | 34951584 | 0.45 | cone | [60] |
| Toxoplasma gondii | Euk | Plunge | none | | 2 | 6 | | 9564564 | 3.00 | capsule | [86] |
| Toxoplasma gondii | Euk | Plunge | none | | 2 | 6 | | 9564564 | 3.00 | capsule | [87] |
| U2OS (human osteosarcoma) | Euk | Plunge | FIB-SEM | | 7 | 30 | | 24764273 | 0.60 | cone | [55] |
| U2OS (U2OS-IRE1a-mNG) (human osteosarcoma) | Euk | Plunge | none | | 7 | 30 | | 24764273 | 0.60 | cone | [75] |
| VeroE6 (monkey kidney) | Euk | Plunge | Ultramicrotome | | 7 | 17.5 | | 31127400 | 1.03 | cone | [10] |
| WI38 (human lung fibroblasts) | Euk | Plunge | none | | 7 | 10 | 90 | 37268154 | 1.80 | rectangular based pyramid | [88] |
